# Using rat operant delayed match-to-sample task to identify neural substrates recruited with increased working memory load

**DOI:** 10.1101/2020.06.18.160028

**Authors:** Christina Gobin, Lizhen Wu, Marek Schwendt

## Abstract

The delayed match-to-sample task (DMS) is used to probe working memory (WM) across species. While the involvement of the PFC in this task has been established, limited information exists regarding the recruitment of broader circuitry, especially under the low- versus high-WM load. We sought to address this question by using a variable-delay operant DMS task. Male Sprague-Dawley rats were trained and tested to determine their baseline WM performance across all (0-24s) delays. Next, rats were tested in a single DMS test with either 0s or 24s fixed delay, to assess low-/high-load WM performance. *c-Fos* mRNA expression was quantified within cortical and subcortical regions and correlated with WM performance. High WM load upregulated overall *c-Fos* mRNA expression within the PrL, as well as within a subset of mGlu5+ cells, with load-dependent, local activation of protein kinase C as the proposed underlying molecular mechanism. The PrL activity negatively correlated with choice accuracy during high load WM performance. A broader circuitry, including several subcortical regions, was found to be activated under low and/or high load conditions. These findings highlight the role of mGlu5 and/or PKC dependent signaling within the PrL, and corresponding recruitment of subcortical regions during high-load WM performance.

## Introduction

Working memory (WM) confers the ability to temporally maintain and manipulate information in the absence of relevant sensory input to guide goal-directed behavior (Cowan 2008). In addition, WM contributes to and overlaps with other domains of executive functioning such as attention, cognitive flexibility and inhibitory control. While WM function changes throughout the life span, pronounced WM impairment been characterized in a number of neuropsychiatric disorders including schizophrenia, substance use disorders and ADHD (Chai et al. 2018). The susceptibility of WM to disruptions stems from the fact that it is a memory system with limited capacity, in which stored information undergoes rapid decay unless. Even if the amount of information in WM does not exceed its capacity, it can impose varied demand on WM processing. This is often referred to as ‘WM load’ and can also be described as the amount of information (or the duration that information) that must be held ‘online’ to solve a particular problem. Imaging studies in humans and non-human primates reproducibly showed that the dorsolateral prefrontal cortex (dlPFC) is a key brain structure activated during WM performance, and further that neural activity within this brain region correlates with the WM load (Manoach et al. 1997; Brzezicka et al. 2018; Toepper et al. 2014). The medial prefrontal cortex (mPFC) in rats is a cortical region thought to be analogous to the human and primate dlPFC, in terms of anatomical connections and function (Seamans et al. 2008; Brown and Bowman 2002). Accordingly, a number of studies using diverse tasks showed that activity within the rat mPFC is required for ‘normal’ WM performance (e.g., Jung et al. 1998; Horst and Laubach 2009; Yang et al. 2014). One of the most commonly utilized behavioral tasks to assess WM in both humans and animals is the (variation of) delayed match-to-sample task (DMS; Daniel et al. 2016; Lind et al. 2015). Operant DMS tasks involve presentation of a stimulus, followed by a delay period and a subsequent choice phase, wherein the selection of a matching stimulus is required to obtain a reward. The advantage of this task that the delay period can be easily adjusted to control the difficulty of the task such that increasing the delay imposes greater WM load. The current study adopted a version of the rat operant DMS task that has been previously shown to require activation of the mPFC (Sloan et al. 2006). A more recent study by (Hernandez et al. 2018) utilizing the same DMS task has identified that the activity of subtype 5 metabotropic glutamate receptor (mGlu5) within this brain region is necessary to maintain high choice accuracy. To further highlight the key role of the mPFC, large hippocampal lesions do not have an impact on the DMS task performance (Sloan et al. 2006). And while, in WM tasks that rely more heavily on spatial navigation (such as Y-maze task, or delayed alternation task in a T-maze) recruitment of the hippocampus has been documented (Vorhees and Williams 2014), detailed investigation of hippocampal activity (or a broader circuitry) involved in the operant DMS task has not been conducted. A recent meta-analysis of human functional neuroimaging studies utilizing the DMS task to investigate WM, confirmed the role of the dlPFC, but also implicated a broader circuitry composed of other cortical areas (e.g., premotor and orbitofrontal cortex), as well as subcortical structures (e.g., thalamus, amygdala; Daniel et al. 2016). It also revealed that variations in the DMS task parameters, such as the use of verbal vs. non-verbal stimuli, results in distinct recruitment of WM networks. And finally, this meta-analysis (and other published evidence; e.g., Chen and Desmond 2005) suggests that activity within some, but not all, human WM-related circuits track variations in WM load.

The current study was motivated by the fact that (besides the mPFC) it is currently unknown which neural circuits are recruited during the operant WM task in rats, and how the activity (or the recruitment) of these circuits varies with increasing WM load (delay). To address this knowledge gap, we conducted a brain-wide analysis of *c-Fos* expression (a well-established marker of recent neuronal activity; Gallo et al. 2018; Morgan and Curran 1991) in rats trained in the operant DMS task and tested under fixed low- and high-load WM conditions. Further, this study also explored the relationship between WM performance and neural activity (*c-Fos* mRNA levels) in the selected brain regions. Lastly, in order to characterize candidate neural and molecular substrates of WM, this study analyzed two variables related to mGlu5 receptor activity in the prelimbic cortex (PrL) immediately following the final WM test: *c-Fos* expression in a subset of mGlu5+ neurons, and the activity of mGlu5 partner protein kinase, protein kinase C (PKC). The results of this study are described below.

## Results

### Delayed match-to-sample task performance under variable and fixed delay conditions

All rats were successfully trained in the DMS task reaching the pre-determined testing criterion in 46.65 ± 3.34 days. The DMS task performance (percent correct responses across the variable 0-24s delay set) was evaluated over two consecutive 5-day testing blocks. A repeated measures two-way ANOVA was conducted with Block and Delay as the within subjects factors. We found a main effect of Delay (*F*(6, 108) = 176.6, *p* < .0001; Fig. 1B), but not Block (*F*(1, 18) = .37, n.s.). The number of trials completed did not differ between Block 1 and Block 2 (data not shown). This suggest a relatively stable overall DMS performance that was sensitive to demand (delay). Prior to the final DMS test, rats were subdivided into three groups: home cage control, fixed 0s and fixed 24s delay groups. Two-way ANOVAs were conducted with Delay as the within-subject factor and Group as the between-subjects factor for each Block that revealed no pre-existing between-group differences in baseline DMS responding. There was no effect of Group (*F*(2, 16) = 0.58, n.s.) and no Group x Delay interaction for Block 1 (*F*(12, 96) = 1.06, n.s.). Also, there was no effect of Group (*F*(2, 16) = 0.63, n.s.) and no Group x Delay interaction for Block 2 (*F*(12, 96) = 0.45, n.s.). The DMS task performance during final test was analyzed using unpaired t-tests to compare percent correct responses (Fig. 1C) and the number of trials completed (Fig. 1D) under the conditions of a fixed (0s or 24s) delay. Rats performed significantly worse under the high-load working memory conditions (24s delay), when compared to the low-load group (0s delay; *t*(13) = 13.6, *p* < 0.0001; Fig. 1C). Rats in the 0s delay group completed more trials than rats in the 24s delay group (*t*(13) = 10.42, *p* < 0.0001; Fig. 1D). The omission rate in both groups was very low (0-4 omissions per test: data not shown).

**Figure 1:**
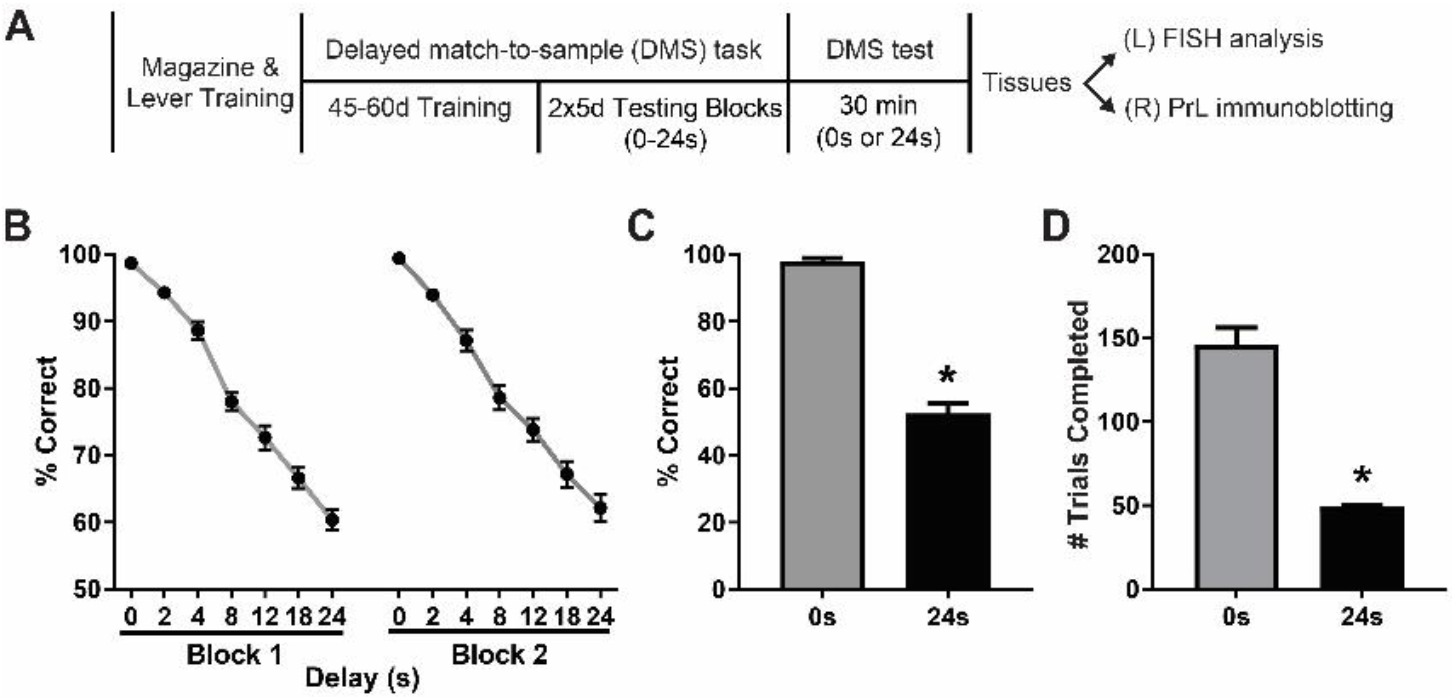
Delayed match-to-sample task performance under a variable, or fixed delay conditions. **A.** Experimental timeline. The left hemisphere (L) was processed for the FISH analysis, while the PrL tissue from the right hemisphere (R) was processed for immunoblotting. **B.** Baseline working memory (WM) testing. Delay-dependent DMS task performance (% correct) under variable 0-24s delay conditions for the period of two consecutive testing blocks. **C-D.** Single DMS test. Rats DMS task performance, % correct responses (C), and the number of trials completed (D) during the single 30 min test with either 0s or 24s fix delay. Mean ± SEM, \**p* < 0.05 vs. 0s group, *n* = 7-8/group.

### *c-Fos* mRNA expression after a fixed delay DMS test

*c-Fos* mRNA levels, used here as a marker of recent neuronal activity, were quantified across several rat brain regions immediately following the final (fixed delay) DMS test, or in age-matched control rats that remained undisturbed in their home cage. *c-Fos* mRNA was quantified as the total mRNA puncta per target region of interest (ROI) averaged from two sections per brain regions for each rat. The ROIs analyzed in the current study have been previously implicated in the regulation of (a) WM performance - prelimbic cortex (PrL), dorsomedial striatum (DmS), nucleus accumbens (NAc), subregions of the hippocampus (CA1 and CA3), perirhinal cortex (PrH), nucleus reuniens (NRe), central amygdala (CeA); (b) - behavioral flexibility (orbitofrontal cortex – OFC); (c) - reward-based decision-making (dorsolateral striatum – DlS), or (d) - served as a positive control for the motor function-related neural activity (primary motor cortex – M1). See the Discussion section for the relevant, region-specific literature references. One-way ANOVAs were used to analyze between-group *c-Fos* mRNA levels (home cage controls, DMS test at 0s and 24s). Significant differences in *c-Fos* mRNA expression were found in the PrL (*F*(2, 17) = 23.10, *p* < 0.0001, Fig. 2A - left), OFC (*F*(2, 16) = 6.89, *p* < 0.01, Fig. 2A - middle), M1 (*F*(2, 16) = 15.45, *p* < 0.001, Fig. 2A - right), DlS (*F*(2, 16) = 4.12, *p* < 0.05, Fig. 2B - left), CA1 (*F*(2, 17) = 5.40, *p* < 0.05, Fig. 2C - left), and NRe (*F*(2, 16) = 7.90, *p* < 0.01, Fig. 2D - left). Tukey’s multiple comparison tests were used for the follow-up analysis of all significant differences. Rats in the 24s condition expressed greater activation (number *c-Fos* mRNA puncta) in the PrL compared to rats in the 0s condition (*p* < 0.01) and home-cage controls (*p* < 0.0001), and rats in the 0s condition expressed more activation in the PrL compared to the home-cage controls (*p* < 0.05, Fig. 2A - left). Compared to home-cage controls, rats in the 0s (*p* < 0.05), and 24s conditions (*p* < 0.01) expressed greater activation (*c-Fos* mRNA puncta) in the OFC (Fig. 2A - middle). Rats in the 0s condition showed greater activation in M1 compared to home-cage controls (*p* < 0.001) and rats in the 24s condition (*p* < 0.01, Fig. 2A - right). Rats in the 24s condition showed greater activation in the DlS compared to home-cage controls (*p* < 0.05, Fig. 2B - left). Compared to home-cage controls, rats in the 0s groups (*p* < 0.05), and 24s conditions (*p* < 0.05) expressed greater activation in CA1 (Fig. 2C - left). Rats in the 24s condition expressed greater activation in the NRe compared to rats in the 0s condition (*p* < 0.05) and home-cage controls (*p* < 0.01, Fig. 2D - right).

**Figure 2:**
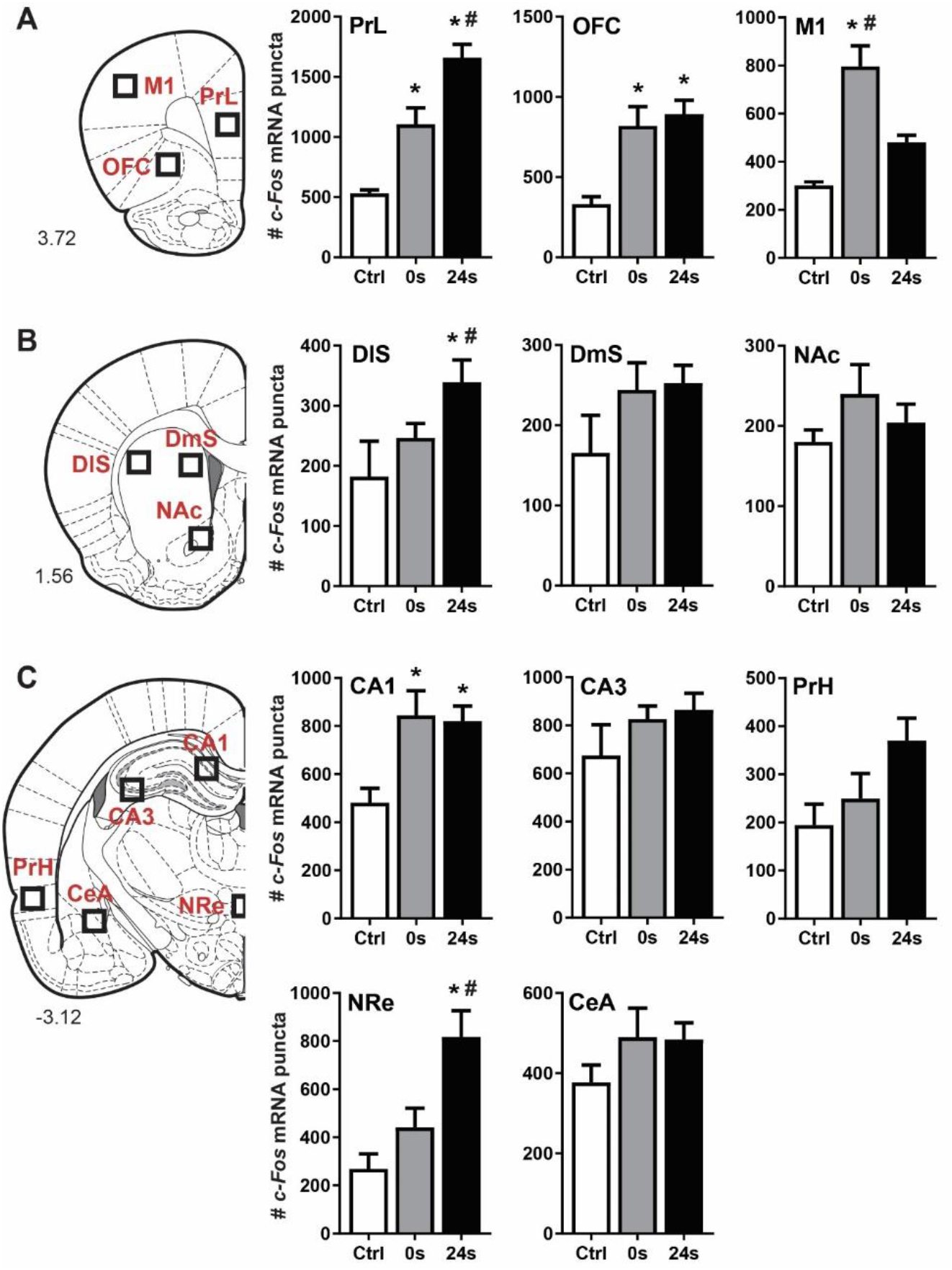
*c-Fos* mRNA expression after a DMS test with fixed 0s and 24s delays. **A, B, C.** (left) Rat brain coronal outlines according to (Paxinos and Watson 2005), with the brain areas used for the *c-Fos* mRNA analysis as highlighted. **A, B, C.** (center and right) Quantitative analysis of *c-Fos* mRNA puncta within the outlines regions of interest corresponding to the prelimbic cortex (PrL), orbitofrontal cortex (OFC), primary motor cortex (M1), dorsolateral striatum (DlS), dorsomedial striatum (DmS), nucleus accumbens (NAc), CA1 and CA3 subregions of the hippocampus, perirhinal cortex (PrH), nucleus reuniens (NRe) and central amygdala (CeA) in rats that underwent a single DMS test with fixed 0s or 24s delays, and in home cage controls (Ctrl). Mean ± SEM, \**p* < 0.05 vs. Ctrl group, #p<0.05 vs. 0s delay group. *n* = 5-8/group.

### Correlations between *c-Fos* mRNA expression and DMS performance at fixed delay test

Bivariate Pearson correlations were conducted to assess the relationship between *c-Fos* mRNA expression (number of puncta) and high-load working memory performance (% correct during the fixed 24s condition). We found a negative correlation between total PrL c-*Fos* mRNA expression and performance at the 24s delay (*r* = −0.72, *n* = 8, *p* < 0.05; Fig. 3B - left). *c-Fos* mRNA expression within the OFC, M1, NRe, CA1, and PrH did not correlate with the performance at the 24s delay (Fig. 3B - middle, right; Fig. 3C).

**Figure 3:**
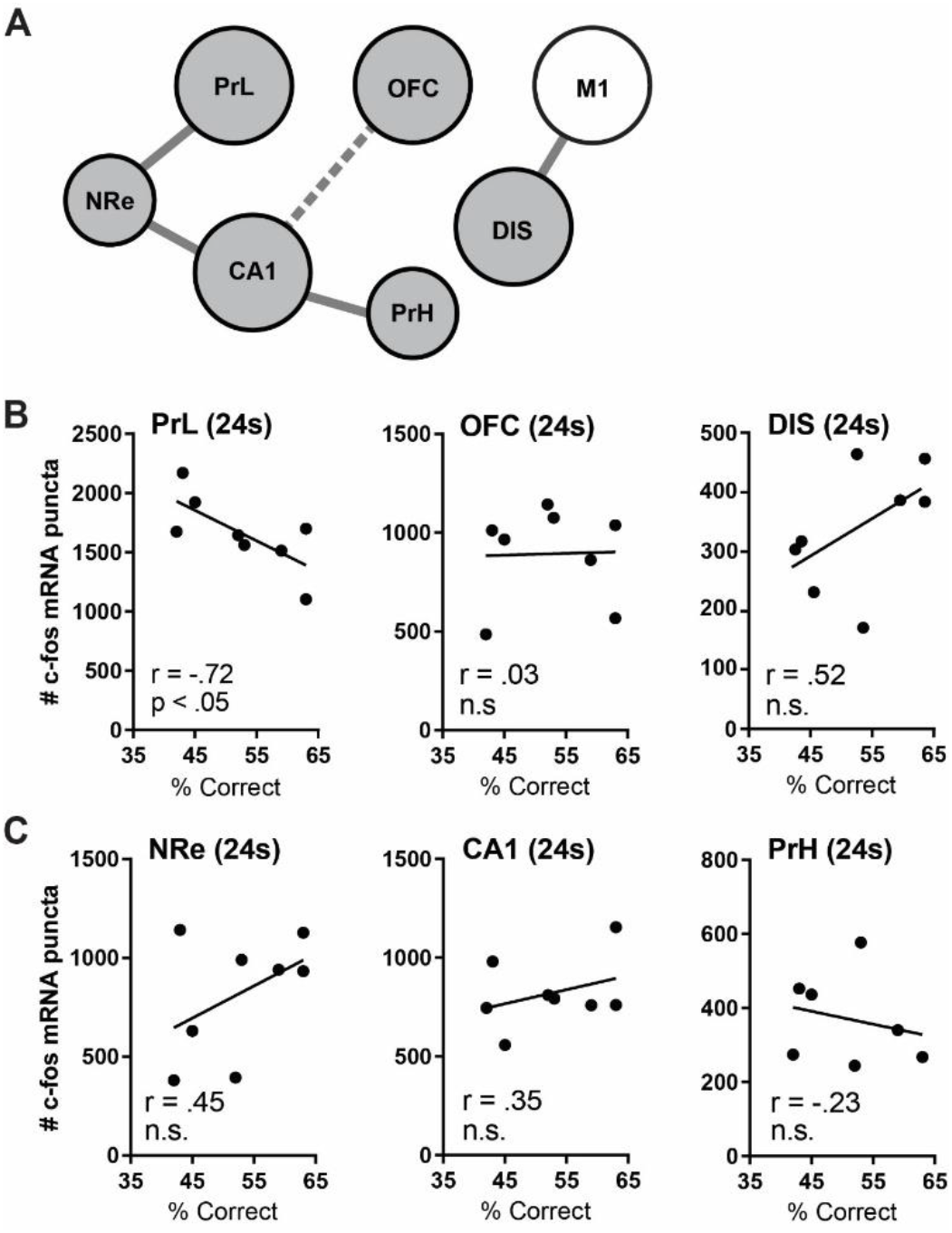
Correlation between *c-Fos* mRNA expression and DMS test performance under a fixed 24s delay conditions. **A.** Predicted neural connections a between the brain regions analyzed. **B-C.** Correlation between the number *c-Fos* mRNA puncta in the PrL, OFC, DlS, NRe, CA1 and CA3 and DMS task performance (% correct responses) under 24s delay condition. *n* = 5-8/group.

### *c-Fos* mRNA expression in mGlu5+ cells in the PrL in relation to fixed delay test DMS performance

The number of *c-Fos* mRNA puncta was averaged across all mGlu5 mRNA expressing cells within the target ROI across two brain sections per rat. A one-way ANOVA was conducted to assess *c-Fos* mRNA expression within PrL mGlu5+ cells between groups and found a significant difference (*F*(2, 17) = 27.32, *p* < 0.0001). Rats in the 24s condition expressed greater activation (average # *c-Fos* mRNA puncta) within PrL mGlu5 expressing cells compared to rats in the 0s condition (*p* < 0.01) and home cage controls (*p* < 0.0001). Rats in the 0s condition expressed greater activation in PrL mGlu5 expressing cells compared to the home cage controls (*p* < 0.05; Fig. 4B - left). Bivariate Pearson correlations were conducted between the average number of *c-Fos* mRNA puncta within mGlu5 expressing cells in the PrL and WM performance (percent correct) during the 0s or 24s conditions. We found a negative correlation between average c-fos expression within mGlu5-expressing cells and DMS performance in the 24s condition (r = −0.79, n =8, p < 0.05, Fig. 4B - middle) but not in the 0s condition (r = 0.04, n.s., Fig. 4B - right).

**Figure 4:**
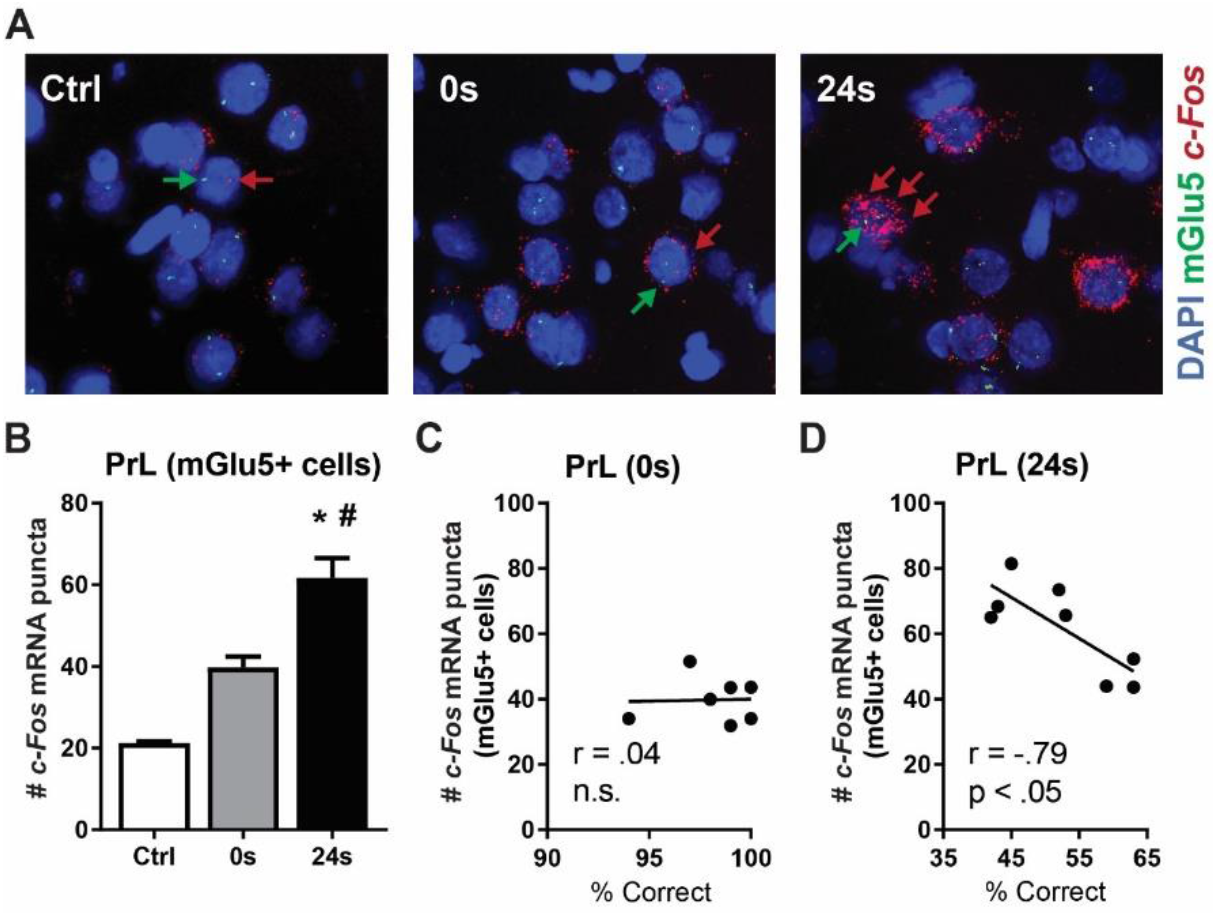
*c-Fos* mRNA expression in mGlu5+ cells in the PrL after a DMS test with fixed 0s and 24s delays. **A.** Representative images of c-Fos mRNA in mGlu5-expressing cells within the PrL for home-cage controls (left), rats in the 0s condition (middle), and rats in the 24s condition (right). *c-Fos* mRNA and *mGlu5* mRNA puncta are indicated in red and green, respectively. Nuclei are counterstained with DAPI (blue). **B.** Quantitative analysis of the *c-Fos* mRNA puncta within the subpopulation of mGlu5+ cells in the PrL in control rats (Ctrl) and in rats that underwent a single DMS test under a fixed 0s or 24s conditions. **C-D.** Correlation between the number of *c-Fos* mRNA puncta and DMS task performance (% correct) under a fixed 0s and 24s delay conditions. Mean ± SEM, \**p* < 0.05 vs. Ctrl group, #p<0.05 vs. 0s delay group. *n* = 5-8/group.

### PKC activity and protein expression in the PrL after a fixed delay DMS test

To further explore possible neurobiological mechanisms recruited during WM task in the PrL, PKC activity in this brain region was evaluated using phospho-(Ser) PKC substrate antibody. This antibody selectively detects phosphorylation state of PKC consensus sites on many cellular proteins, providing an indirect measure of tissue PKC activity (Chopra et al.; Kim et al. 2010; Bilodeau and Schwendt 2016). Due to a different signal intensity, PKC-phosphorylated proteins with molecular weight over 120kDa (high kDa) were analyzed separately from the <120kDa proteins (low kDa). A one-way ANOVA revealed significant group differences in PKC activity (PKC substrate phosphorylation) for both high kDA (*F*(2, 17) = 4.28, *p* < 0.05) and low kDA (*F*(2,17) = 4.01, *p* <0.05) groups of proteins (Fig. 5B – left and middle). Follow-up post-hoc analysis showed increased PKC-mediated phosphorylation of high kDA (*p* < 0.05) and low kDA (*p* < 0.05) substrates in the PrL of rats tested under 24s delay conditions. Though trend in increased PKC phosphorylation of low kDa substrates was observed at 0s delay (p = 0.07; Fig. 5B). No relationship between the magnitude of PKC substrate phosphorylation in the PrL and high load working memory performance (% correct at the 24s delay condition) was found (data not shown). In contrast to varied PKC activity, no group differences in the overall PKC content in the PrL were found (Fig. 5B – right). Immunoblotting analysis also did not find group differences in c-Fos protein content (Fig. 5C) and mGlu5 dimer and monomer levels in the PrL (Fig. 5D – left and right). As the duration of the final DMS test (30 min) was too short to alter protein levels, this data indicate that there were no preexisting group differences in c-Fos, mGlu5 or PKC protein content in the PrL.

**Figure 5:**
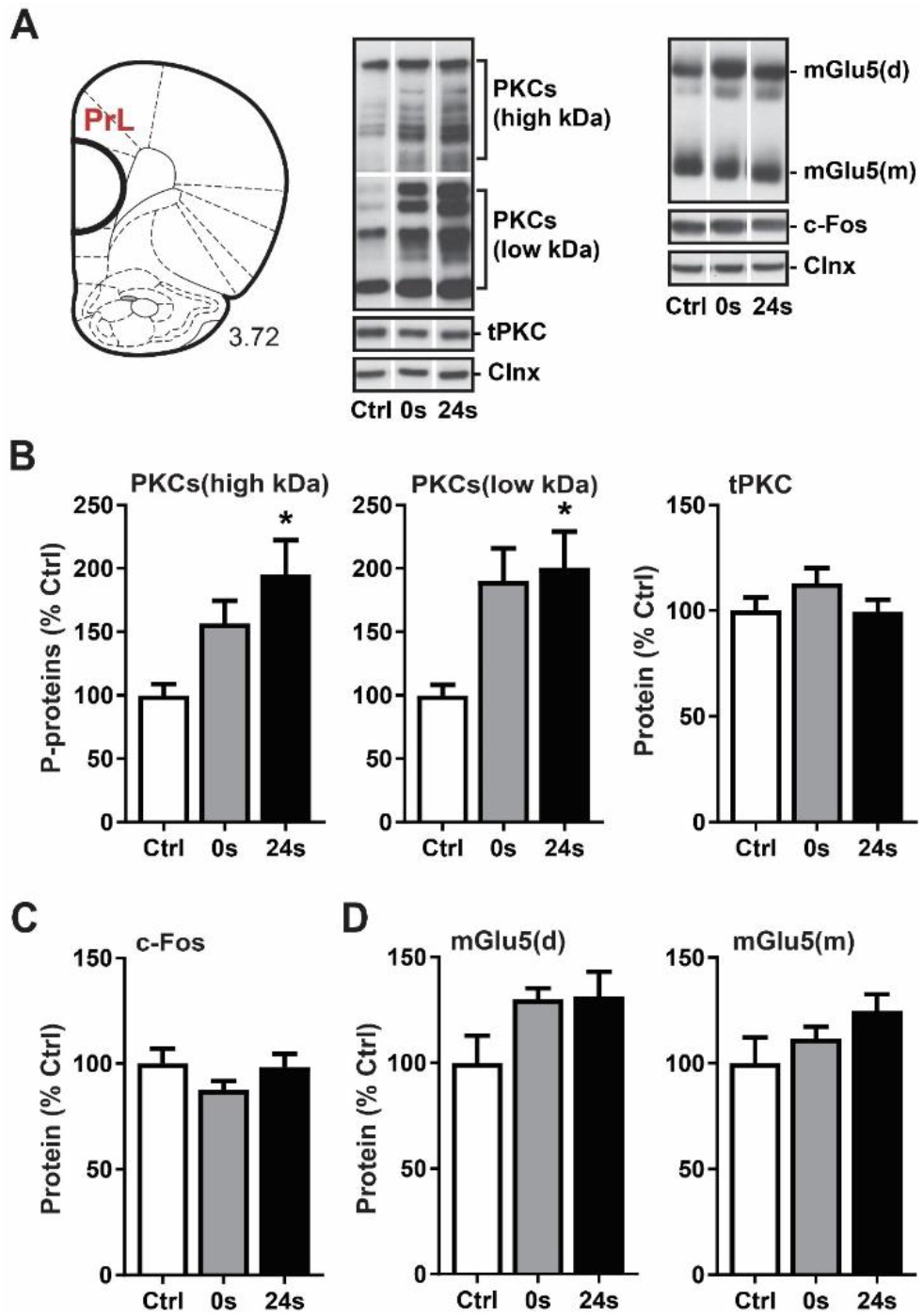
PKC activity and protein expression in the PrL after a DMS test with fixed 0s and 24s delays. **A.** (left) A rat brain coronal outline according to (Paxinos and Watson 2005), with the PrL dissection site highlighted. (right) Representative immunoreactive bands as detected with antibodies against phospho-(Ser) PKC substrates (PKCs), total PKC (tPKC), c-Fos protein, mGlu5 receptor (monomer and dimer) and loading control, calnexin (Clnx). Numbers to the left of each blot correspond to molecular weight of proteins in kDa. **B-D.** Quantitative immunoblotting analysis of high- / low-molecular weight phospho-PKC substrates and tPKC, c-Fos protein and mGlu5 monomer and dimer. Mean ± SEM, \**p* < 0.05 vs. Ctrl group. *n* = 5-8/group.

## Discussion

The current study characterizes *c-Fos*-expressing neuronal populations recruited with an increasing WM load in rats performing an operant DMS task. It identifies the PrL as the brain region, most sensitive to variations in WM load, with a broader circuitry that includes dorsolateral striatum (DlS), nucleus Reuniens (NRe), and the CA1 subregion of the hippocampus activated under low and/or high load conditions. On the other hand, neuronal activity detected in the primary motor cortex (M1) reflects motor activity, not the task difficulty. This study also found that under the conditions of high WM load, *c-Fos* mRNA levels in the PrL negatively correlate with the performance accuracy in the DMS task, suggesting inefficiency in neuronal processing. Initial exploration of the neural and molecular substrates activated with an increased WM load revealed that increasing WM load (a) upregulates *c-Fos* expression in a subset of PrL neurons that also express mGlu5 and (b) is associated with hyperactivity of PKC, the main intracellular kinase associated with this receptor. This indicates that the PrL (and some other directly or indirectly connected brain areas) are recruited with increasing WM load and that the magnitude of mGlu5- and/or PKC-dependent signaling in this brain region predicts WM performance under these conditions. The present findings also complement our recent research showing that chronic cocaine impairs DMS task performance in rats, particularly under the increased WM load conditions, and that aberrant PrL activity (assessed via immediate-early gene expression) is related to both post-cocaine WM deficits and persistent cocaine-seeking (Hámor et al. 2020; Gobin et al. 2019).

Here, we utilized an operant DMS task that allows for repeated, programmed trial-by-trial changes in WM load. Under these conditions, the choice accuracy (percent correct choices per session) ranged from ~ 98% (at 0s delay) to ~ 62% (at 24s delay) during the baseline DMS testing. We also observed that the overall baseline DMS task performance is relatively stable in well-trained rats as no differences between testing Block 1 and 2 were detected. However, once the task parameters were altered in the final test (changed from a random presentation of seven different 0-24s delays to a fixed 0s or 24s delay) performance in the 24s group suffered and declined to near-chance levels. Rats in the 24s group also completed significantly fewer trials compared to their 0s counterparts. This is indicative of very demanding task conditions under which rats possibly experienced high reward-delivery uncertainty and ‘frustration’ that is known to negatively affects WM performance in humans (Fillauer et al. 2020). Even though ‘frustration’ is an anthropomorphic concept, some evidence suggests that ‘frustration effect’ can account for the altered the strength of reinforced and non-reinforced responses in rats (Stout et al. 2003).

The main goal of this study was to follow-up on the WM performance data with an assessment of neural activity, measuring *c-Fos* mRNA expression. Specifically, we utilized this approach to quantify neural activity/*c-Fos* expression across a number of brain regions at the end of the single DMS test conducted under fixed 0s or 24s delay conditions, corresponding to low and high WM load, respectively. Out of all brain regions analyzed, the PrL displayed a unique pattern of *c-Fos* mRNA expression that mirrored increasing WM load. This was a novel, though not surprising finding, as the PrL is a key regulator of rodent WM (Vertes 2004; Arime and Akiyama 2017). Persistent firing and synchronization within PrL neurons have been identified during the delay period in a WM task and implicated in the encoding and representation of WM (Yang et al. 2014; Jung et al. 1998; Constantinidis et al. 2018). Consequently, acute inactivation, lesioning, or pathological changes within the PrL result in WM deficits (Sloan et al. 2006; Izaki et al. 2001; Arime and Akiyama 2017) and others. This is also true for the type of operant DMS task used in this study, as lesioning the mPFC (including the PrL) severely impairs WM performance in this task (Sloan et al. 2006). Interestingly, mPFC lesions did not impair the ability to adapt to rule switching (from match-to-sample to non-match-to-sample; Sloan et al. 2006), indicating that other brain regions (such as the OFC; discussed below) mediate cognitive flexibility in this task.

Beyond the PrL, two other cortical areas analyzed also showed altered *c-Fos* mRNA levels after the final DMS test. In the OFC, increased *c-Fos* mRNA levels were detected in all rats exposed to the final DMS test regardless of the delay used. This finding is in agreement with the OFC-centric circuits controlling cognitive flexibility, rather than WM performance (Fettes et al. 2017; Barbey et al. 2011). It could be hypothesized that switching from a variable 0-24s delay to a fixed (0s or 24s) delay, calls for an update in the learned behavioral strategies, which in turn requires bringing the OFC online. Accordingly, activation of this brain region occurs in response to changes in the DMS task parameters (from variable to fix delay, both DMS groups), not increased task difficulty (delay, 24s group only). Research in human subjects suggests that the OFC is involved in the coordination of multiple WM processes in situations when the application of prior cognitive strategies is not sufficient to achieve a behavioral goal. In support, patients with damage to the OFC display deficits in complex tasks that require coordination of WM maintenance, manipulation, and monitoring processes, but not in simple tests of WM maintenance (Barbey et al. 2011). In contrast to both the PrL and the OFC regions, *c-Fos* expression in the primary motor cortex (M1) reflected recent motor activity, with rats in the 0s test group showing significantly upregulated *c-Fos* mRNA and the most trials completed, in comparison to 24s group (fewer trials completed) or home cage controls (rest, no trials).

We have also evaluated a number of subcortical regions. In the striatum, increased *c-Fos* mRNA levels were detected in the DlS under the 24s delay conditions, but not in the DmS or NAc. The lack of *c-Fos* changes in the DmS was somewhat surprising, as prior studies have indicated the involvement of this brain region in controlling the choice accuracy in operant and T-maze-based delayed non-match to position (DNMTP) tasks (Akhlaghpour et al. 2016; Li et al. 2018). It should be noted that our results do not rule out the role for cortico-striatal projections in WM performance; instead, they suggest that depending on task rules (DMS vs. DNMTP; novel vs. well-trained), or altered frequency of reward delivery, cortico-striatal map might include the DlS not DmS (Balleine et al. 2007). Alternatively, as firing pattern in the DmS is transient and shifts throughout the delay period (unlike the persistent firing of the delay cells in the PrL; Akhlaghpour et al. 2016), DmS neuronal activity might not be sufficient to generate measurable differences in *c-Fos* mRNA levels.

The midline thalamic nucleus termed the NRe has been assigned the role of an important brain structure supporting reciprocal hippocampal-mPFC communication. It is believed that the NRe relays and regulates spatial and contextual information between these brain structures, contributing to the synchronized firing of the mPFC and hippocampus under conditions of increased WM load (for reviews see: Griffin 2015; Dolleman-van der Weel et al. 2019). In agreement, our data confirm a delay-dependent activation of the NRe (increased *c-Fos* mRNA levels in 24s delay group). This suggests that NRe is recruited even during WM tasks with a lesser spatial component, as compared to spatial WM tasks, such as delayed alternation task in a T-maze (Viena et al. 2018). The NRe neurons that receive input from mPFC send dense excitatory projection to the CA1 subregion of the dorsal hippocampus (Wouterlood et al. 1990; Herkenham 1978). It is this projection that is responsible for the synchronous cortico-hippocampal activity during memory processing and consolidation (Hauer et al. 2019). Here we found an increase in *c-Fos* mRNA levels in the CA1 (but not in the CA3) subregion of rats after the final DMS test, regardless of the length of the delay imposed. It is possible that in addition to the PrL, other cortical regions (perhaps the OFC) interact with the CA1 to offer behavioral flexibility under changed experimental conditions, such as when variable delay testing is replaced by the single delay in the final DMS test. Even though not well understood, the interactions between the OFC and the hippocampus is thought to promote cognitive flexibility during learning, memory, and decision making (Wikenheiser and Schoenbaum 2016), suggesting that analogous pattern of *c-Fos* mRNA upregulation detected in the OFC and the CA1 is not accidental. The role of CA1 in supporting continuous DMS task performance should be further evaluated, as studies disagree whether the global hippocampal lesions impair WM in operant DMS/DNMS tasks (Sloan et al. 2006; Broersen 2000; Aggleton et al. 1992). To broaden the interpretation of the CA1 *c-Fos* data, we analyzed neuronal activity related to the recent DMS task performance in the PrH. PrH is a cortical region that has robust reciprocal connections with the hippocampal formation, in particular with the CA1 subregion of the hippocampus (Kealy and Commins 2011). By controlling the information flow into and out of the CA1, the PrH has been shown to participate in cognitive processes, such as object- or stimulus-recognition memory (Lee and Park 2013). However, limited evidence suggests that selective lesions of the PrH produced deficits in an operant delayed non-matching-to-position task in rats, that were more pronounced at longer delays (Wiig and Bilkey 1994; Wiig and Burwell 1998). Here, we failed to find a significant upregulation of c-Fos mRNA in the PrH, though a trend towards an increase was detected at the longer 24s delay. Finally, to assess the specificity of the observed patterns of neuronal activation for the DMS task, we evaluated *c-Fos* mRNA in the CeA, a brain nucleus linked to long-term emotional memory, but not necessary for immediate WM (Bianchin et al. 1999). In agreement, we found no differences in *c-Fos* mRNA in the CeA.

Beyond uncovering region-specific and demand-dependent differences in neuronal activation during WM testing, we sought to investigate the relationship between choice accuracy (percent correct responses) and *c-Fos* mRNA levels. Out of the six brain regions in which *c-Fos* mRNA was elevated under the high load (24s delay condition), only the PrL *c-Fos* levels correlated with choice accuracy. As discussed above, a switch from the variable 0-24s delay set (baseline testing) to a forced 24s delay in the final DMS test, increases task difficulty. It is possible that the observed negative correlation between the PrL *c-Fos* levels and high load WM performance reflects (a) recruitment of additional network capacity within the PrL to support new learning required to improve task performance and increase the frequency of reward delivery, or (b) it is a non-specific upregulation of the PrL activity due to ‘frustration effect’ related to inability to receive reward.

Since the PrL activity showed the highest sensitivity to WM load, this study set out to further characterize neural and molecular substrates within this brain region activated during WM testing. While the neural mechanisms of WM are complex (for reviews see: (Arnsten and Jin 2014; Goldman-Rakic 1995), studies in humans and animals suggest that glutamatergic activity in the Prl (or dlPFC) is necessary for ‘normal’ WM performance. In this regard, a recent study by (Woodcock et al. 2018) showed that glutamate levels rise in the human dlPFC in response to increased WM demand. On the other hand, inhibiting glutamate release via local administration of mGlu2/3 (auto)receptor agonists, or blockade of the postsynaptic ionotropic glutamate receptors disrupted sustained firing of local pyramidal neurons and impaired WM performance in animals (Gregory et al. 2003; van Vugt et al. 2020; Wang et al. 2013). Recent studies by our laboratory and others, showed that systemic or local intra-PrL inhibition of mGlu5 receptors impairs DMS task performance (Gobin and Schwendt 2020; Hámor et al. 2020; Hernandez et al. 2018). This study reveals for the first time that increasing WM demand progressively recruits mGlu5-positive cells in the PrL as evidenced by increased *c-Fos/mGlu5+* mRNA in the 24s (but not in the 0s) delay group. Similar to overall *c-Fos* expression, *c-Fos/mGlu5+* expression in the PrL negatively correlates with choice accuracy, suggesting that activity within this neuronal subpopulation serves to compensate or overcome behavioral inefficiency.

Finally, as PKC is the major cellular effector downstream from mGlu5 receptors implicated in WM (Dash et al. 2007; Runyan et al. 2005; Birnbaum et al. 2004b), this study analyzed whether the activity of this kinase in the PrL tracks increased WM load (delay). The phosphorylation status of peptide sequences selectively recognized by the PKC served as a measure of PKC-mediated cellular phosphorylation, and ultimately, as an accurate indicator PKC tissue activity (Bilodeau and Schwendt 2016; Chopra et al.; Kim et al. 2010). We found that phosphorylation of both high- and low-molecular-weight PKC substrates was increased in the 24s delay group. This corresponds with findings of increased PKC activity in the mPFC of unimpaired rats immediately after a delayed-match-to-place test in a Morris water maze (Runyan et al. 2005). As PKC is crucial for memory formation in many other brain regions (for review see: (Sun and Alkon 2014), the question arises, whether the observed increase in the PrL PKC activity is promoting, or degrading WM. Previously, excessive activation of PKC in the mPFC has coincided with the stress-induced disruption of WM, while in aged rats, PKC activity predicted WM impairment (Brennan et al. 2009; Birnbaum et al. 2004b). And further, systemic or intra-PFC inhibition of PKC improved WM performance in normal young rats, or rescue stress- or aging-related WM impairment (Birnbaum et al. 2004; Hains et al. 2009; Brennan et al. 2009). The molecular mechanism of how abnormal PKC activity disrupts WM performance is not clear, but can include interference with the delay-related activity of the dedicated pyramidal neurons (Birnbaum et al. 2004a). This can explain how elevated PKC in the 24s delay group can contribute to poor WM performance observed in our study. However, as WM requires ‘optimal’ cognitive processing, and both hypofunction or hyperactivity of delay cells can negatively affect WM, future studies should investigate the possibility that the relationship between PKC activity and WM is more complex, perhaps following an inverted U-shape curve, akin to the effects of increasing dopaminergic and glutamatergic tone in the PFC on WM (Jin et al. 2017; Cools and D’Esposito 2011; Vijayraghavan et al. 2007).

## Materials and Methods

### Animals

Adult male Sprague-Dawley rats (Charles River Laboratories; 275g on arrival; N = 20) were first acclimated to the animal facility prior to any manipulation. They were housed individually, maintained on a 12 hr reverse light/dark cycle (lights off at 0700), and given *ad libitum* access to water, and food-restricted (15-20 g of food per day) to maintain ~ 85% of their free-feeding weight as previously described (Gobin & Schwendt, 2017; Gobin & Schwendt, 2019). All animal procedures were approved by the Institutional Animal Care and Use Committee of the University of Florida and performed in accordance with the Guide for the Care and Use of Laboratory Animals. The overall experimental timeline is depicted in Figure 1A.

### Operant Delayed Match-to-Sample Task: training and testing under a variable delay condition

Rats were trained and tested in the delayed match-to-sample (DMS) task, as previously described (Gobin & Schwendt, 2019). Briefly, rats were subjected to a daily 40 min training (or testing) sessions (one session per day) in standard rat operant chambers (30 × 24 × 30 cm; Med Associates, St. Albans, VT) equipped with two levers. Each session began with an illumination of the house light, which remained on throughout the session, except during time-outs. Each trial consisted of three phases: a sample phase, a delay period and a choice phase. In the sample phase, a left or right lever was randomly selected by the computer such that there was equal presentation of each lever throughout the session. Pressing the sample lever resulted in retraction of that lever, delivery of a sucrose pellet, and initiation of the delay interval with randomized delay durations. During the choice phase, both levers were presented. Pressing the lever previously presented during the sample phase, resulted in delivery of a sucrose pellet and the correct response was recorded. Pressing the other lever was scored as an incorrect response and resulted in a time-out period wherein no sucrose pellet was delivered, the house light was extinguished, and both levers were retracted for a duration of 6s prior to the start of the next trial (time-out period). First rats underwent a 30 min magazine training session wherein 29 sucrose pellets were delivered into the food hopper at random intervals, and rats were required to consume all of these pellets prior to progressing to lever press shaping. During lever press shaping, rats were presented with only the left lever in a 60 min session. Upon reaching a criterion of at least 50 lever presses on the left lever (each rewarded with a sucrose pellet), they underwent lever press shaping for the right lever the following day with the same criterion. During the next training phase, rats were trained without any delays between the sample and choice phase, including a correction procedure to prevent the development of side biases. Next, rats were trained at two delay sets: short delay set {0s, 1s, 2s, 3s, 4s, 5s, 6s}, and intermediate delay set {0s, 2s, 4s, 8s, 12s, 16s}. Rats were required to reach a criterion of ≥ 80% correct responses over two consecutive days prior to progressing to the next delay set. Finally, upon reaching criterion at the intermediate training delay set, rats entered the testing phase for two blocks of five days at the final delay set {0 s, 2s, 4s, 8s, 12s, 18s, 24s}.

### Operant Delayed Match-to-Sample Task: the final test under a fixed delay conditions

Following the completion of the two DMS testing blocks, rats were assigned to either 0s or 24s test (or home cage control) condition and subjected to a final DMS test. This test was modified to last 30 min (corresponding to a peak in stimulus-induced *c-Fos* response; Gallo et al. 2018), and each trial was limited to a presentation of a single delay (0s or 24s) to test DMS performance under low-load (easy), or high-load (hard) WM conditions (n = 7-8/group). Controls rats (n = 5) were handled but remained in their home cage.

### Tissue Collection and Processing

Rats were euthanized by decapitation immediately following the final DMS test (or after their removal from the home cage). Rat brains were rapidly extracted, left and right hemispheres separated, and snap-frozen in isopentane (2-methylbutane) chilled on dry ice. The right hemisphere was cut into serial 12μm coronal brain sections using a cryostat (Leica CM1950). Tissue sections were collected as follows: PrL, M1, and OFC were collected approximately at +3.72, DlS, DmS, and NAc were collected at approximately +1.56, while sections containing CA1, CA3, PrH, CeA, and NRe were collected approximately at −3.12 relative to Bregma (Paxinos & Watson, 2005). Sections were freeze-mounted onto Superfrost Plus Gold slides (Fisher Scientific), air-dried, and stored at −80°C. The PrL tissue from the left hemisphere was collected using a 2 mm micropunch (Harris Uni-Core, Ted Pella, Redding, CA) as shown on Fig. 5A. All tissues were stored at −80°C for later fluorescent in situ hybridization (right hemisphere sections), or western blotting analysis (left hemisphere PrL punches).

### Fluorescent in situ Hybridization

Fluorescent in situ hybridization (FISH) for c-*Fos* and *mGlu5 (Grm5)* mRNAs was performed using standard or custom-designed validated target riboprobes (Gobin et al. 2019), ACDBio) and RNAscope Multiplex Fluorescent Reagent Kit (ACDBio, Newark, CA) following published procedures (Wang et al., 2012; Gobin et al., 2019). Briefly, rat brain sections were fixed in 4% paraformaldehyde (PFA) for 15 min at 4℃, dehydrated in a series of graded EtOH concentrations for 5 min each (50%, 70%, 100%, 100%) and dried at room temperature prior to undergoing 25-min protease digestion using pretreatment #4 (ACDBio). RNAscope target probes for *c-Fos* (ACDBio: 403591-C1, lot 18179C) and *mGlu5* (*GRM5*; ACDBio: 471241-C2, lot 17243B) were applied to each section, and slides were incubated at 40°C for 2hr. Next, preamplifier and amplifier probes were applied to each section and incubated at 40°C (AMP 1, 30 min; AMP 2, 15 min; AMP 3, 30 min). The AMP 4 Alt-C was selected so that *c-Fos* and *mGlu5* probes were labeled with ATTO 550 and Alexa 647 fluorophores, respectively. Sections were counterstained with DAPI and coverslipped with ProLong Gold antifade mounting reagent (Thermofisher). Fluorescent images were obtained using an Olympus BX51 microscope equipped with an apochromatic 40x objective, CCD high-resolution camera, and ISIcapture software (both by Tucsen Photonics, Fuzhou, China). Quantification of mRNA puncta was performed in ImageJ and MATLAB (Mathworks, Natick, MA) using TransQuant software (Bahar Halpern and Itzkovitz 2016). Cells expressing both *c-Fos* and *mGlu5* mRNA transcripts were identified manually with ImageJ (Schneider et al., 2012). Segmentation of all dual *c-Fos*/*mGlu5*-positive cells was performed in TransQuant and within-cell *c-Fos* mRNA puncta were measured.

### Immunoblotting

The tissue protein lysates were prepared from the PrL as described previously (Bilodeau and Schwendt 2016). Total protein content was quantified using bicinchoninic acid (BCA) assay according to the manufacturer’s instructions (BCA Protein Assay Kit, Thermo-Fisher, Waltham, MA). Equal amounts of total protein (15 μg/lane) were separated by SDS-PAGE (4–15% polyacrylamide) and transferred onto polyvinylidene difluoride (PVDF) membranes. Membranes were blocked for one hour in 5% milk in Tris-buffered saline buffer that included 0.1% Tween 20 (TBST), and probed overnight with the following primary antibodies: rabbit anti-phospho-(Ser) PKC substrates (1:15,000; Cell Signaling #2261), mouse anti-pan PKC (1:10,000, MilliporeSigma #05-983), rabbit anti-c-FOS (1:10,000; MilliporeSigma, #ABE457,), and rabbit anti-mGlu5 (1:5,000; MilliporeSigma, #AB5675). PKC substrate antibody selectively detects proteins with phosphorylated PKC consensus sites (phosphorylated Ser residues surrounded by Arg or Lys at the –2 and +2 positions and a hydrophobic residue at the +1 position, Cell Signaling, Danvers, MA). Next, membranes were washed three times in 5% milk/TBST and incubated with a species-matched HRP-conjugated secondary antibody (1:20,000, Jackson Immuno Research, West Grove, PA). Immunoreactive bands on the membranes were visualized with a chemiluminescent reagent (ECL+) using a high performance chemiluminescence film (both from: GE Healthcare, Piscataway, NJ). Equal loading and transfer of proteins was verified with a reversible protein stain (Ponceaus S) and by re-probing membranes with a housekeeping protein calnexin (1:20,000, #ADI-SPA-860, Enzo Life Sciences, Farmingdale, NY, USA). Integrated band density of each protein sample was measured using Image Studio Lite software (LI-COR Biosciences, Lincoln, NE, USA), normalized it to its respective calnexin integrated density measure, and expressed as the percentage of home-cage controls values.

### Statistical Analysis

GraphPad Prism (Version 8.4.2) software was used to analyze all data with the alpha level set at 0.05 for all statistical analyses. Repeated measures two-way ANOVA was used to compare within-subject variables (Delay and Block) during DMS testing. One-way ANOVAs and t-tests were used to compare group differences on single dependent measures. Tukey’s multiple comparison tests were used to follow up significant effects. Two-tailed Pearson correlations were conducted to compare continuous variables.

## Acknowledgments

The authors thank Dr. Jennifer Bizon and Dr. Joseph McQuail for their input on the design of the DMS task, as well as Peter Hámor and Lori Knackstedt for their valuable comments on the manuscript. We also thank Mr. Spencer Berman, Mr. James Hickman, and Mr. Jason Dee for their assistance with behavioral studies. This research was supported by the University of Florida McKnight Brain Institute Pilot Grant awarded to MS.

